# A neuropeptide signaling system that rapidly enforces paternity in the *Aedes aegypti* mosquito

**DOI:** 10.1101/136150

**Authors:** Laura B. Duvall, Nipun S. Basrur, Henrik Molina, Conor J. McMeniman, Leslie B. Vosshall

## Abstract

Female Dengue and Zika vector mosquitoes (*Aedes aegypti)* generally mate once, with sperm from this male fertilizing all eggs produced in her lifetime. Here we implicate HP-I, an *Aedes*- and male-specific neuropeptide transferred to females, and its cognate receptor in the female, NPYLR1, in rapid enforcement of paternity. *HP-I* mutant males were ineffective in enforcing paternity when a second male was given access to the female within 1 hour. *NPYLR1* mutant females produced mixed paternity offspring at high frequency. Synthetic HP-I injected into wild-type virgins reduced successful matings, but had no effect on *NPYLR1* mutant females. Asian tiger mosquito (*Ae. albopictus*) HP-I potently activated *Ae. aegypti* NPYLR1. Invasive *Ae. albopictus* males are known to copulate with and sterilize *Ae. aegypti* females, and cross-species transfer of HP-I may contribute to this phenomenon. This neuropeptide system promotes rapid paternity enforcement within *Ae. aegypti*, but may promote local extinction in areas where they compete with *Ae. albopictus*.

**One Sentence Summary:** *Aedes*-specific peptide rapidly enforces paternity

**Text:** *Ae. aegypti* females typically mate only once with one male in their lifetime, a behavior known as “monandry” (1). This single mating event provisions the female with sufficient sperm to fertilize the >500 eggs she will produce during her ∼4-6 week lifespan in the laboratory (2). Successful mating is capable of inducing lifetime refractoriness to subsequent insemination by other males, enforcing the paternity of the first male (3-5). In other species, males use diverse strategies to assure the paternity of their offspring, for instance physical barriers such as mating plugs found in mice (6) and *Anopheline* mosquitoes (7), and anti-aphrodisiac pheromones used by *Drosophila melanogaster* males to tag female flies as non-virgin (8). Another widely used strategy in insects is the transfer of biologically active male seminal proteins, produced by the male accessory gland and secreted into the ejaculatory duct along with sperm during insemination, to affect the sexual receptivity of the female (3, 9-13). Perhaps the best-characterized male seminal fluid protein in insects is the *Drosophila* fly sex peptide (11), which acts on the sex peptide receptor in the female to suppress receptivity and trigger egg production (12). *Drosophila* sex peptide receptor mutant females will readily remate with multiple males, and wild-type females that mate with sex peptide mutant males remain sexually receptive.

---

*Ae. aegypti* mate in flight near human hosts (14). Males and females modulate their wing beat frequency in a mating duet (15, 16), but the role of this auditory communication in female choice is not completely understood. Copulation is extremely rapid, lasting less than 15 seconds (4, 17). Remarkably, females become refractory to remating within seconds, bending their abdomen to deter additional males from successful copulation (1, 3, 4). This suggests that there must be a rapid mechanism to enforce the paternity of the first male, by preventing female re-mating. Previous work has identified protein components in male seminal fluid that induce long-term female sexual refractoriness with a slow onset similar to that seen for sex peptide in *Drosophila* (3). But the signals that induce short-term refractoriness remain unknown.

In the course of investigating a role for Head Peptide-I [HP-I; so named because of its initial detection in mosquito heads (18)], in female host-seeking behavior (19), we discovered that HP-I is a factor that exerts enforcement of paternity within one hour of mating. *Ae. aegypti* HP-I is a member of the short neuropeptide F (sNPF) family, and probably resulted from the duplication of the sNPF gene in the *Aedes* lineage (20). Based on extensive BLAST searches of genome and short-read databases, HP-I appears to be unique to *Aedes* species. It has a close homologue in *Ae. albopictus*, but no known homologues in other species (Fig. 1A). The *HP-I* gene encodes a pre-pro-peptide (Fig. 1C) that is post-translationally modified to produce three identical copies of HP-I, which are then further modified by the addition of hydroxy-proline and c-terminal amidation to produce mature HP-I (Fig. 1D) (18).

**Fig. 1.**
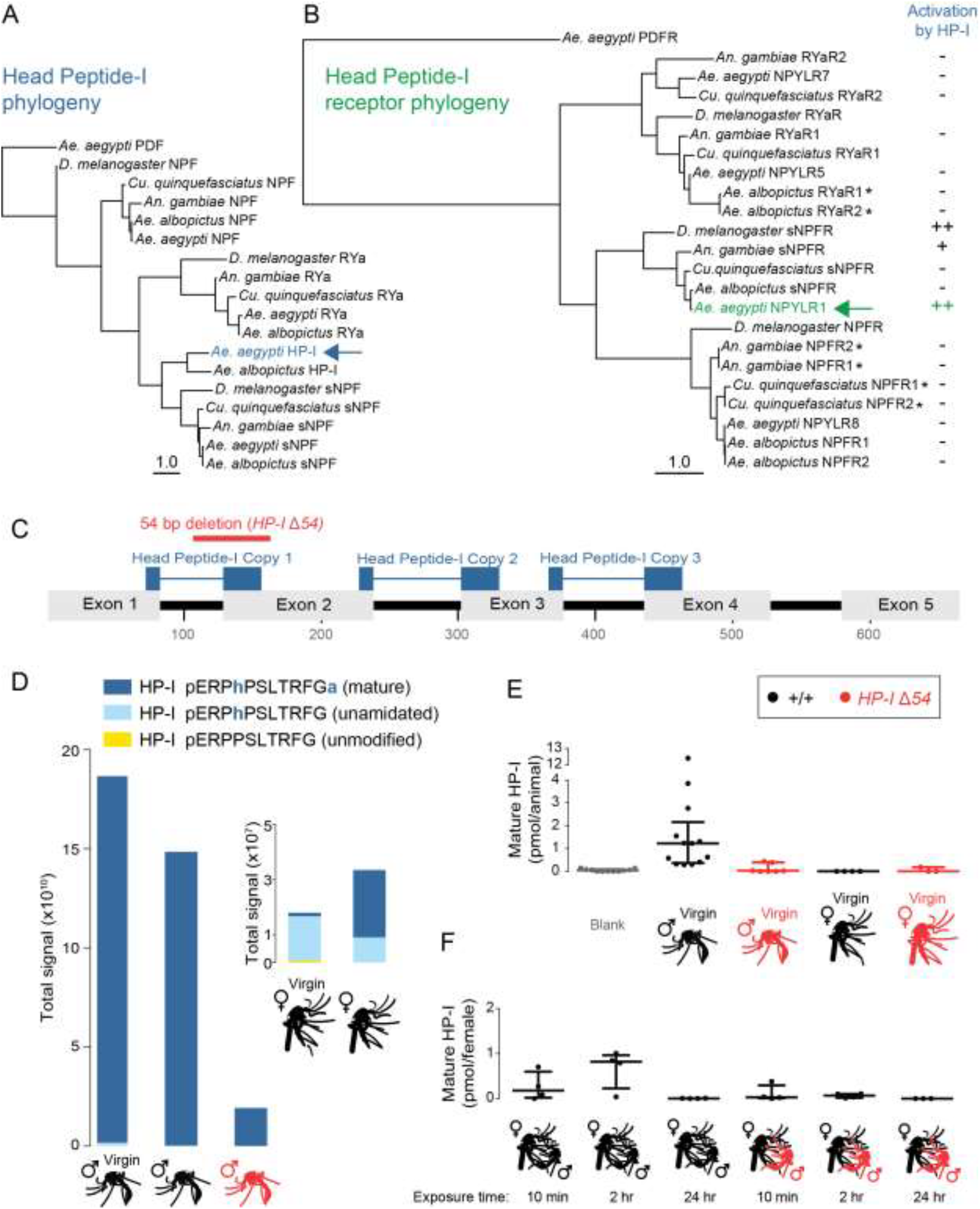
HP-I is an *Aedes*-specific, male-enriched neuropeptide. (**A-B**) Phylogenetic protein trees of HP-I and related neuropeptides (pre-pro-peptides) (A) and NPYLR1 and related neuropeptide receptors (B), with PDF and PDFR as the outgroup, respectively. Branch lengths represent mean expected rate of amino acid substitution. Scale = 1.0. Pairs of receptors marked with an asterisk encode highly similar or identical proteins that are annotated as separate genes. Column at right indicates relative activation of indicated receptors by 50 μM *Ae. aegypti* HP-I in cell-based assays. (**C**) Structure of the *HP-I* gene and the position of the *HP-IΔ54* mutation. (**D**) Relative levels of mature and immature HP-I detected by LC-MS in whole adult mosquitoes of the indicated sex, genotype, and mating status with females shown as inset due to dramatically lower levels of HP-I (n = 1) (**E**) Quantification of mature HP-I detected by LC-MS/MS in whole adult mosquitoes of the indicated sex and genotype (n = 4-13 groups of 7-10 animals). (**F**) Quantification of mature HP-I detected by LC-MS/MS in whole adult female mosquitoes of the indicated genotype who were exposed to males of the indicated genotype for the indicated time (n = 4 groups of 7-10 females). Data in E-F are shown as median with interquartile range.

Our previous work identified neuropeptide Y-like receptor 1 (NPYLR1) as the HP-I receptor in *Ae. aegypti* (21). NPYLR1 is a member of the insect sNPF receptor family (Fig. 1B). To examine the specificity of HP-I, we carried out cell-based assays with three major classes of neuropeptide receptors, RYaR, sNPFR, and NPFR, from 5 insect species (Data File S1). We found that *Ae. aegypti* NPYLR1 was strongly activated by HP-I, and that HP-I did not activate RYa receptors or NPF receptors in any of these species (Fig. 1B). HP-I activated more distantly related sNPFRs in *D. melanogaster* and *An. gambiae*. To test the *in vivo* function of HP-I, we used CRISPR-Cas9 (22) to generate a 54 bp deletion spanning intron 1 and exon 2, which is predicted to disrupt copy 1 of HP-I and introduce a frame shift (Fig. 1C). These *HP-IΔ54* mutants developed normally and showed no gross developmental delays or abnormalities (data not shown).

Although HP-I levels were previously reported to increase in females following a blood-meal (19), later studies did not detect HP-I in female tissues (23). Naccarati et al. (24) showed that the male accessory gland is the primary source of the peptide. We used liquid chromatography mass spectrometry (LC-MS) to confirm these results and to characterize HP-I levels in the *HP-I* mutant. Mature HP-I was detected at very high levels in wild-type males, greatly reduced levels in mutant males, and was nearly undetectable in wild-type females (Fig. 1D). The presence of residual HP-I in the mutant suggests that it is a hypomorph, perhaps due to repair of the frameshift by splicing to downstream exons. This extreme sexual dimorphism was not observed for 4 other neuropeptides also detected in all samples (Data File S1).

Naccarati et al. (24) showed that HP-I is transferred from males to females, where it is detectable for only ∼2 hours after mating. We carried out quantitative LC-MS/MS to measure levels of mature HP-I in wild-type and *HP-I* mutant virgin males and females (Fig. 1E), and wild-type virgin females exposed to wild-type or *HP-I* mutant males for varying periods of time (Fig. 1F). Wild-type males produced high levels of mature HP-I, mutant males produced lower levels, and mature HP-I was nearly undetectable in both wild-type and *HP-I* mutant females (Fig. 1E). We detected high levels of HP-I in wild-type females exposed to wild-type males for 10 minutes and 2 hours, but not 24 hours (Fig. 1F). There were nearly undetectable levels of HP-I in females exposed to *HP-I* mutant males for any amount of time.

Since males are the source of HP-I that is transferred to females, we examined blood-feeding behavior and fecundity in females mated to *HP-I* mutant males. Females blood-fed to the same extent (Fig. 2A and B), consumed blood-meals of the same size, produced eggs with the same temporal profile (Fig. 2C) in the same quantity (Fig. 2D), regardless of their genotype or that of the males they were exposed to. This suggests that HP-I transferred from the male during mating plays no significant role in controlling egg production, in strong contrast to the sex peptide system in *Drosophila* (11, 12).

**Fig. 2.**
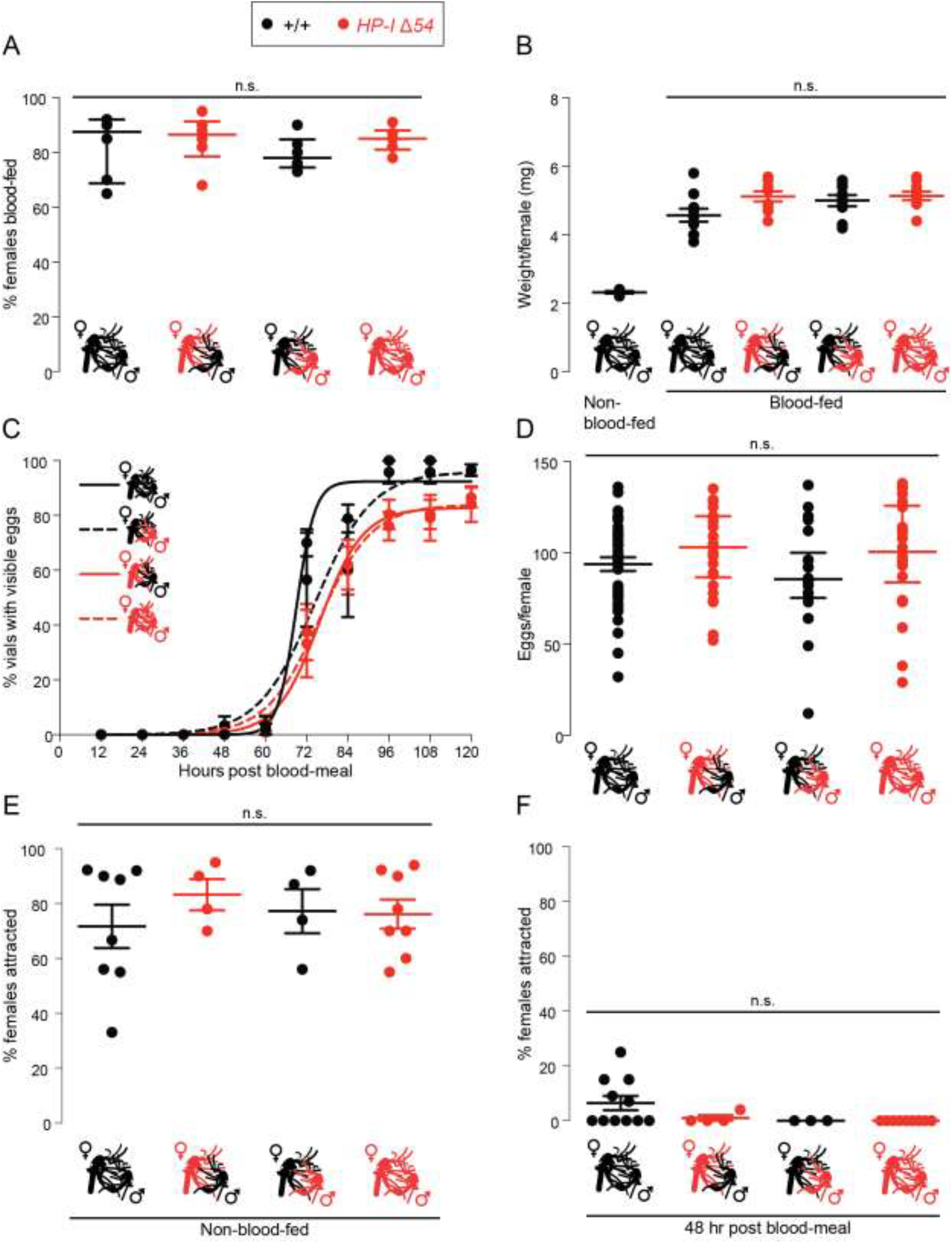
HP-I mutant females show normal reproductive and host-seeking behavior. (**A**) Blood-feeding success of females of the indicated genotype who were exposed to males of the indicated genotype for >24 hours (median with interquartile range; n = 6 trials, 25 - 40 females per trial; Kruskal-Wallis test, n.s. = not significant). (**B**) Weight of fully-blood-fed females in (A) immediately after blood-meal (data points represent the average weight per animal, weighed in groups of 10 females; mean ± SEM, n=5-11 groups of 10; ANOVA followed by post-hoc Bonferroni correction for multiple comparisons, n.s. = not significant). (**C**) Temporal profile of female egg-laying of the indicated genotype exposed to males of the indicated genotypes for >24 hours (mean ± SEM; n = 5 trials, 12-24 females per trial). There was no significant difference in between the genotypes at any given time point (p>0.05 Kruskal-Wallis test). (**D**) Eggs laid per individual female in (C) (mean ± SEM; n = 22-39; ANOVA followed by Bonferroni correction for multiple comparisons, n.s. = not significant). (**E-F**) Host attraction of females of the indicated genotypes, exposed to males of the indicated genotype for >24 hours, before (E) and 48 hours after (F) a blood-meal (mean ± SEM n = 3 – 11 trials, 15 - 20 females per trial; Kruskal-Wallis test n.s. = not significant).

HP-I was previously thought to mediate suppression of host-seeking behavior after a blood-meal (19, 21), either by being produced in females in response to a blood-meal (19) or by being transferred from the male during mating (24). We therefore measured attraction to human host cues of wild-type and *HP-I* mutant females exposed to wild-type or *HP-I* mutant males, before and 48 hours after a blood-meal. We found no effect of the *HP-I* mutation on attraction to human hosts before a blood meal (Fig. 2E) or suppression of host attraction 48 hours after a blood-meal (Fig. 2F). These findings are consistent with our previous work showing that the HP-I receptor *NPYLR1* is not required for female mosquito fecundity, host-seeking, blood-feeding, or egg-laying behaviors (21).

Given its enrichment in males and transfer to females during mating, we investigated a role for HP-I in mating success. *HP-I* mutant males were equally capable of fathering offspring when mated to wild-type females, as compared to wild-type males or a paternity marker strain carrying a transgenic marker with ubiquitous expression of ECFP (Fig. 3A). Over half of females successfully produced offspring after only 5 minutes of exposure to any of the three genotypes of males, with nearly 100% success after 30 minutes of exposure (Fig. 3A). These results reinforce the observation that mating occurs very rapidly in *Ae. aegypti* (1, 3, 4).

**Fig. 3.**
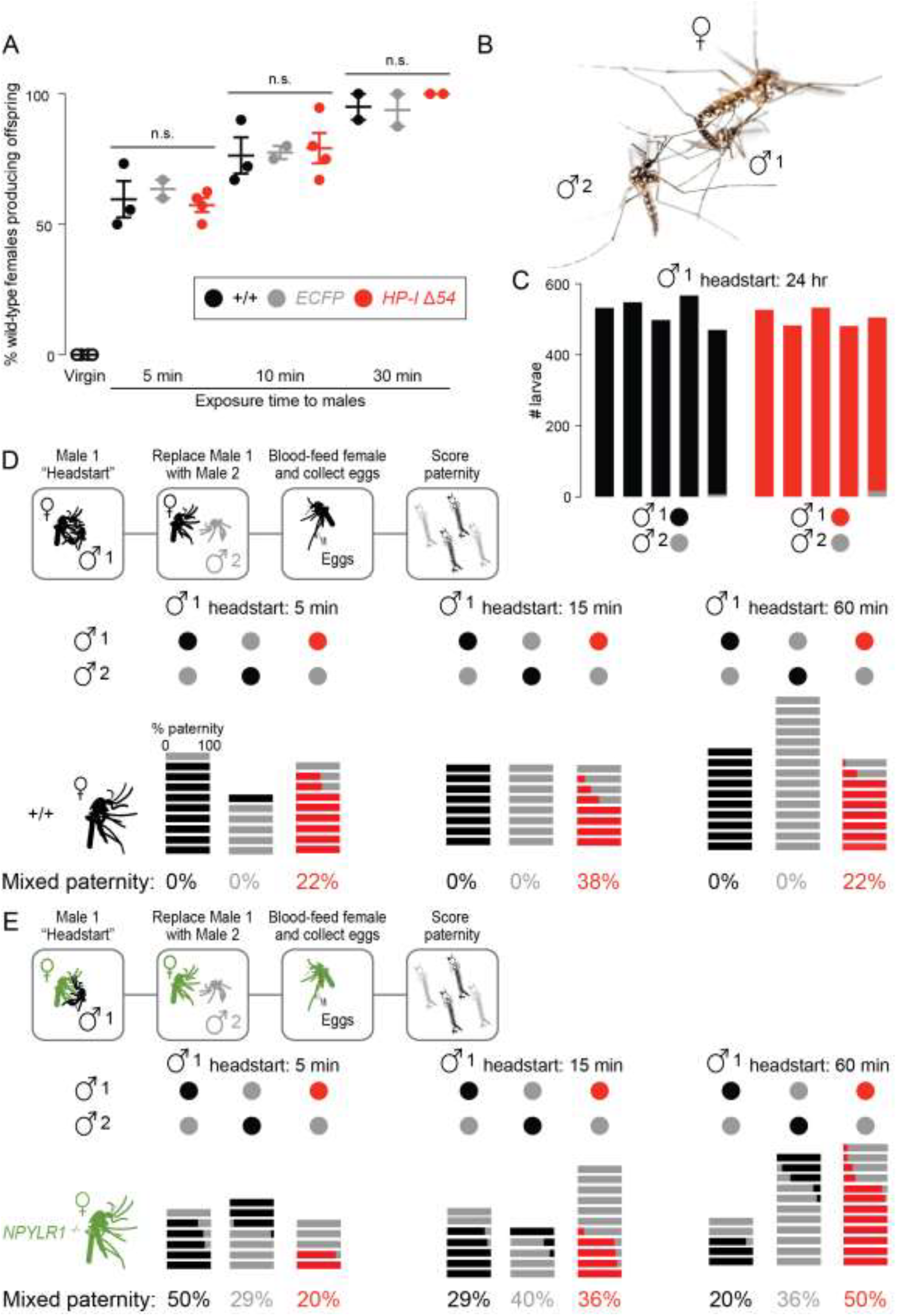
*HP-I* mutant males mate normally but fail to enforce paternity. (**A**) Offspring produced by wild-type virgins or females exposed to males of the indicated genotype for the indicated time (mean ± SEM, n = 2 - 4 trials, 18 - 24 females per trial. Kruskal-Wallis test, n.s. = not significant). (**B**) Aerial mating of wild-type *Ae. aegypti*. Photo: Alex Wild. (**C**) Paternity enforcement in groups of 6 wild-type females exposed to 7 males of the indicated genotype for 24 hours (n = 5 trials, 6 females per trial). Data indicate the paternity of offspring. (**D-E**) Top: Schematic of paternity experiments with wild-type (D) or *NPYLR1* mutant (E) females. Bottom: Paternity enforcement in individual wild-type females (n = 6 - 15) (D) or *NPYLR1* mutant females (n = 5 - 12) (E) exposed to males of the indicated genotype for the indicated time. Each horizontal bar represents the offspring of a single female colored to indicate the paternity of her offspring. Mixed paternity of each group is indicated at the bottom.

Female *Ae. aegypti* are generally monandrous (1, 3), which poses the question of how remating is rapidly suppressed, and how the paternity of the first male is enforced (Fig. 3B). We used remating assays in which groups of females were sequentially exposed to males of two different genotypes, and the paternity of the offspring determined. We defined paternity enforcement as the ability of a male to father all offspring produced by a female despite her subsequent exposure to other potential mates. Previous work reported levels of remating in the field (25) and in semifield conditions as high as 14% (26). It is important to note that these earlier experiments used different methods to establish paternity, either microsatellite analysis of offspring (25) or detection of labeled seminal fluid derived from two different males (26). Our laboratory work directly scored the genotype of each live offspring using the ECFP paternity marker strain.

In these experiments, offspring of groups of 6 females were pooled and scored for paternity Offspring fathered by wild-type or *HP-I* mutant males were non-fluorescent, and easily distinguished from their ECFP-positive half-siblings. *HP-I* mutant males, given a 24 hour head start, showed no deficit in enforcing their paternity (Fig. 3C). This is consistent with our quantitative LC-MS/MS experiment showing that HP-I is a short-acting factor in females, not detectible 24 hours after mating (Fig. 1F) (24). It further suggests that additional male factors transferred along with HP-I enforce paternity on the scale of days to weeks (9). In a single trial, we observed a very small number of ECFP-positive offspring among the much larger number of wild-type or *HP-I* mutant offspring. We cannot distinguish between the possibilities that this was a rare polyandrous event or that a female mated only with ECFP-positive male 2.

We next asked whether HP-I is required to enforce paternity within one hour of exposure to the male. In these experiments, offspring of single females were scored individually. This method allowed us to attribute any mixed paternity offspring to acceptance of multiple mates by a single female. When male 1 was wild-type or carried the ECFP marker, there was no mixed paternity whether the exposure time was 5, 15, or 60 minutes (Fig. 3D). In experiments with wild-type and ECFP males given a 5 minute headstart, we occasionally observed offspring fathered exclusively by male 2. We assume that these are instances where male 1 did not mate with the female during the head start he was offered (see Fig. 3A). In contrast to the monandry enforced by wild-type males, *HP-I* mutant males failed to enforce their paternity, and we observed offspring fathered by both males at all time points tested (Fig. 3D). Because the *HP-IΔ54* mutation is a hypomorph, it is conceivable that an HP-I protein null mutant would show more complete failure to enforce paternity. Since NPYLR1 is the only known HP-I receptor in *Ae. aegypti* (Fig. 1B) (21), we asked if *NPYLR1* is required in females to enforce male paternity. Indeed, offspring of *NPYLR1* mutant females (21) showed mixed paternity at all time points tested, regardless of the genotype of male 1 or male 2 (Fig. 3E).

If HP-I transferred from a male to a female during mating enforces his paternity, HP-I injected directly into a virgin female should interfere with subsequent mating success. To test this, individual virgin females were injected with buffer, mature HP-I, or control peptides, and allowed to recover for 12-16 hours in groups. This extended recovery time was required because virgins tested shortly after injection did not mate regardless of the substance injected (data not shown). After recovery, injected females were exposed to wild-type males for 30 minutes, an exposure time that was sufficient for nearly all females to produce offspring (Fig. 3A). >70% of females injected with buffer, inactive HP-I (Cys-10), and two other neuropeptides successfully produced offspring. In contrast, <40% of females injected with mature HP-I produced offspring (Fig. 4A). These results are consistent with the idea that HP-I acts as a signal to the female that she has already mated. We note that suppression of successful mating following HP-I injection was not complete, perhaps because the dose of injected HP-I was lower than that after mating. If the HP-I receptor NPYLR1 mediates the detection of male-derived HP-I, then *NPYLR1* mutant virgins should be insensitive to HP-I injection. Although the overall success of offspring production was lower in *NPYLR1* mutants injected with any peptide, HP-I had no significant effect on success in producing offspring compared to control peptide and buffer injections (Fig. 4B).

**Fig. 4.**
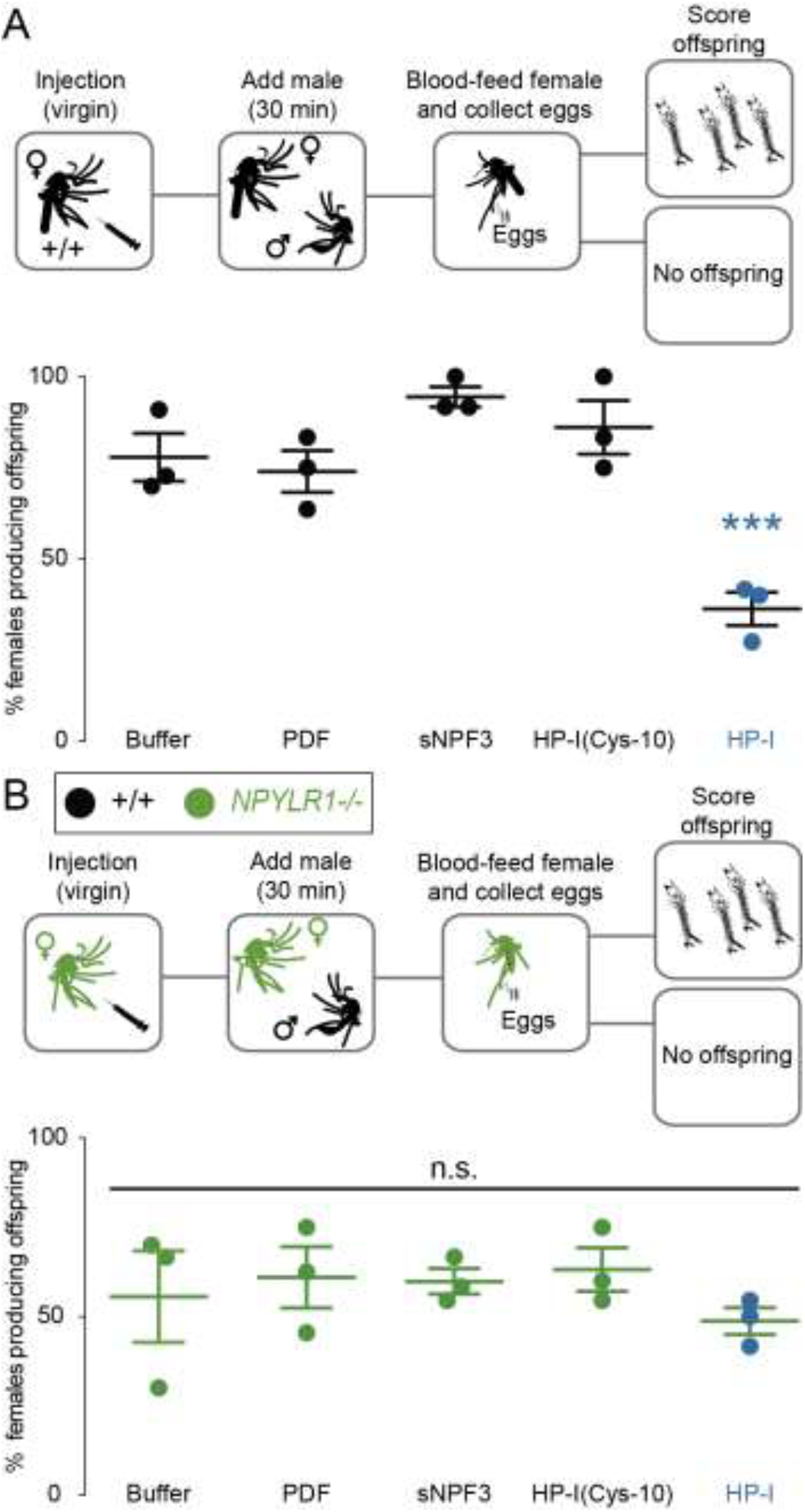
Injected HP-I interferes with reproduction in wild-type but not *NPYLR1* mutant females. (**A**) Top: schematic of wild-type female injection experiments. Bottom: production of offspring by females injected with the indicated peptides (mean ± SEM, n = 3 trials with 5-12 wild-type females per trial. ANOVA with Bonferroni correction, ***p < 0.001). (**B**) Top: schematic of *NPYLR1* mutant female injection experiments. Bottom: production of offspring by females injected with the indicated peptides (mean ± SEM, n = 3 trials with 5-12 *NPYLR1* mutant females per trial. ANOVA with Bonferroni correction, n.s. = not significant).

Since the 1980s, invasive Asian tiger mosquitoes (*Ae. albopictus*) have been displacing *Ae. aegypti* throughout the southern United States, and field observations have documented instances of *Ae. aegypti* females inseminated by *Ae. albopictu*s males (27). Since no offspring are generated by these pairings, this cross-species mating effectively sterilizes *Ae. aegypti* females by preventing subsequent mating with *Ae. aegypti* males. This phenomenon was first observed by Phil Lounibos and colleagues, who termed it satyrization (27) (right panel, Fig. 5A).

**Fig 5:**
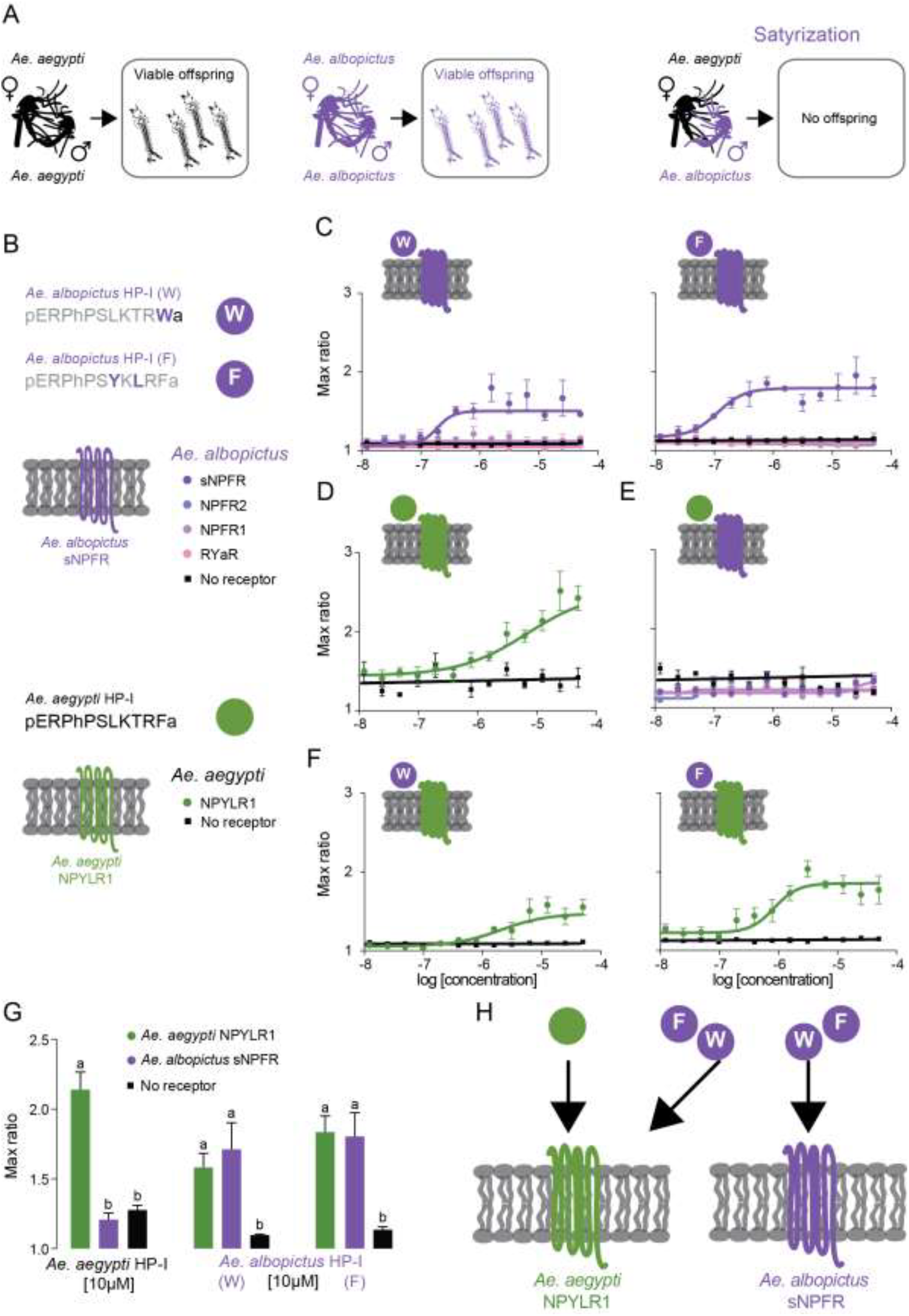
*Ae. albopictus* HP-I peptides are potent activators of *Ae. aegypti* NPYLR1. (**A**) Schematic of normal within-species mating (left and middle) and cross-species satyrization of *Ae. aegypti* females by *Ae. albopictus* males (right). (**B**) Legend for cell-based assay experiments in C-G, including the amino acid sequences of predicted mature HP-I in *Ae. albopictus* compared to *Ae. aegypti* HP-I. (**C**) Dose-response curves of *Ae. albopictus* HP-I peptides (W and F) on *Ae. albopictus* neuropeptide receptors. (**D**) Dose-response curve of *Ae. aegypti* HP-I on *Ae. aegypti* NPYLR1. (**E**) Dose-response curves of *Ae. aegypti* HP-I on *Ae. albopictus* neuropeptide receptors. (**F**) Dose-response curves of *Ae. albopictus* HP-I peptides (W and F) on *Ae. aegypti* NPYLR1. (**G**) Responses to 10μM *Ae. aegypti* and *Ae. albopictus* HP-I peptides. Data in C-G are shown as max ratio (maximum fluorescence level/baseline fluorescence level) (mean ± SEM, 3 replicates. *** p < 0.0001, 1-way ANOVA followed by Tukey’s multiple comparisons test). (H) Schematic of activity of *Ae. aegypti* and *Ae. albopictus* HP-I peptides against *Ae. aegypti* NPYLR1 and *Ae. albopictus* sNPFR.

Importantly, satyrization is unidirectional. *Ae. aegypti* males do not sterilize *Ae. albopictus* females (28). It is possible that the seminal fluid proteins that promote monandry within a species are capable of acting across species to promote satyrization. If this is the case, *Ae. albopictus* male-derived proteins should activate signaling pathways in *Ae. aegypti* females, but the cognate proteins in *Ae. aegypti* should fail to activate *Ae. albopictus* pathways.

The *Ae. albopictus* HP-I gene (Fig. 1A) is predicted to produce two different peptides. Of these, one inferred mature peptide is identical to *Ae. aegypti* HP-I with the exception of a terminal RW-amide instead of RF-amide. The second predicted *Ae. albopictus* HP-I mature peptide has a terminal RF-amide, but two internal substitutions compared to *Ae. aegypti* HP-I (Fig. 5B). We used HEK293T cell-based assays to express candidate HP-I receptors from both mosquito species and monitored receptor activation by these peptide ligands. Based on its homology to *Ae. aegypti* NPYLR1, we predicted that *Ae. albopictus* sNPFR would be the HP-I receptor (Fig. 1A). Indeed, both *Ae. albopictus* HP-I (W) and *Ae. albopictus* HP-I (F) activated *Ae. albopictus* sNPFR, but not *Ae. albopictus* receptors from the NPF and RYa family (Fig. 5C). Consistent with our earlier published work (21), *Ae. aegypti* HP-I activated *Ae. aegypti* NPYLR1(Fig. 5D). Although *Ae. aegypti* HP-I is a potent agonist of the HP-I receptor in its own species, this peptide did not activate any of the *Ae. albopictus* receptors, including the *Ae. albopictus* HP-I receptor, sNPFR (Fig. 5E). While there is no cross-species activation of the *Ae. albopictus* receptor by the *Ae. aegypti* peptide, both of the *Ae. albopictus* HP-I peptides strongly activated the *Ae. aegypti* HP-I receptor, NPYLR1 (Fig. 5F and G). This unidirectional cross-species activity suggests that *Ae. albopictus* male-derived peptides may show biological activity in *Ae. aegypti* females. The transfer of these peptides during cross-species mating could contribute to satyrization by *Ae. albopictus* males by activating the HP-I neuropeptide system in *Ae. aegypti* females and inappropriately enforcing the “paternity” of the *Ae. albopictus* male (Fig. 5H).

In *Ae. aegypti* mosquitoes, mating occurs rapidly in the context of the human host with competing males in close proximity (14), and it is advantageous for males to quickly induce their mate to reject future suitors. Our results show that HP-I is a major mechanism by which paternity is enforced shortly after mating. Long-term monandry is HP-I-independent, and likely relies on other male seminal proteins (3, 5, 10, 29). The mechanisms by which rapid-acting HP-I works with other longer-acting male seminal proteins to enforce paternity remain to be discovered. It will be important to determine in what cells the HP-I receptor, NPYLR1, is expressed in the female. The *Drosophila* sex peptide receptor is expressed in sensory neurons in the female reproductive tract that project to the abdominal ganglion of the ventral nerve cord (30). These neurons relay mating information to central circuits to trigger long-term changes in sexual receptivity (31). The sex peptide system exerts its control over female reproduction over a relatively long time-scale, starting between 4-12 hours after mating (11, 12), and is not permanent. Wild-type *Drosophila* females will remate with additional males within 5-10 days of initial mating (32, 33) Because the *Ae. aegypti* HP-I system acts more rapidly than the sex peptide system, it may use a local circuit within the female mosquito reproductive tract to sense HP-I in seminal fluid and switch off receptivity on a rapid time scale.

Monandry can be exploited by other species, as suggested by the observation that *Ae. albopictus* can satyrize *Ae. aegypti* females. Our discovery that *Ae. albopictus* HP-I can activate *Ae. aegypti* NPYLR1 *in vitro* suggests a mechanism by which this signaling system contributes to cross-species mating interference. It is intriguing to note that just as *Ae. aegypti* males cannot satyrize *Ae. albopictus* females, *Ae. aegypti* HP-I does not activate the *Ae. albopictus* HP-I receptor.

Because HP-I is unlikely to be the sole factor that enforces lifetime monandry in either species, it is likely that additional long-acting factors transferred from *Ae. albopictus* males are biologically active in *Ae. aegypti* females. Both of these species are invasive disease vectors that pose an increasing threat to public health. The release of genetically modified sterile males (34), relies on monandry to be effective because sterile males father no offspring, but they still make females refractory to subsequent mating. This reduces disease transmission by reducing vector mosquito populations. Further exploration of the mechanisms of paternity enforcement will enable more effective application of genetic control strategies, and better understanding of the natural dynamics of interspecies competition.

## Acknowledgments

We thank Laura Harrington, Kevin Lee, Jeff Liesch, Laura Kramer, Phil Lounibos, Lindy McBride, Mariana Wolfner, Nilay Yapici, and members of the Vosshall Lab for discussion and comments on the manuscript. Ben Matthews and Molly Liu helped generate the *HP-I*Δ*54* mutant strain. Gloria Gordon and Libby Mejia provided expert assistance with mosquito rearing. Stable isotope HP-I was synthesized by Henry Zebroski. This research was supported in part by grants from the National Institutes of Health (UL1RR024143 to The Rockefeller University, R01 DC014247 to L.B.V.). L.B.D. was supported by a Rockefeller University Women & Science Fellowship, and by a Postdoctoral Fellowship from the American Philosophical Society. L.B.V. is an investigator of the Howard Hughes Medical Institute.

## Author contributions

L.B.D. carried out all experiments with the exception of the LC-MS/MS experiments in Fig. 1, which were carried out in collaboration with H.M., and the HP-I injections in Fig. 4, which were carried out by N.B. C.J.M. generated the ECFP paternity marker strain. L.B.D. and L.B.V. designed the experiments, interpreted the results, and with the other co-authors composed the figures and wrote the paper.

The authors declare no conflicts of interest.

All raw data reported in this paper are presented in Data File S1.

## Supplementary Materials for

**This file includes:**

Materials and Methods

**Other Supplementary Material for this manuscript includes the following:**

Data File S1

## Materials and Methods

### Mosquito rearing and maintenance

*Aedes aegypti* wild-type laboratory strains (Orlando) were maintained and reared at 25-28°C, 70-80% relative humidity with a photoperiod of 14 hours light:10 hours dark (lights on at 7 a.m.) as previously described (35). Adult mosquitoes were provided constant access to 10% sucrose. Adult females were blood-fed on mice for stock maintenance, on human subjects for HP-I mutant generation, and on human subjects or sheep blood delivered via Glytube membrane feeders (36) for egg-laying and host-seeking experiments. Female mosquitoes were fasted for 14-24 hours in the presence of a water source prior to behavioral experiments. Blood-feeding procedures with live hosts were approved and monitored by The Rockefeller University Institutional Animal Care and Use Committee and Institutional Review Board, protocols 15772 and LV-0652, respectively. Human subjects gave their written informed consent to participate.

### Phylogenetic trees

Predicted protein sequences of pre-pro-peptide and receptor genes were downloaded from VectorBase or UniProt and aligned with MUSCLE (37). Maximum likelihood phylogenetic trees for pre-pro-peptides and receptors were constructed with RaxML (38) using the PROTGAMMAJTT model, testing nodes with the rapid bootstrap analysis (100 replicates). The outgroup was PDF and PDFR for the peptide and receptor tree, respectively. Trees were visualized with Interactive tree of life (39). Peptide genes (accession numbers): *Ae. aegypti* PDF (AAEL001754), *Ae. aegypti* NPF (AAEL002733), *Ae. aegypti* RYa (AAEL011702), *Ae. aegypti* sNPF (AAEL012542), *Ae. albopictus* NPF (AALF017136), *Ae. albopictus* RYa (GAPW01005454.1), *Ae. albopictus* sNPF (LOC109427954), *Ae. albopictus* HP-I (LOC109398455), *An. gambiae* NPF (AGAP004642), *An. gambiae* RYa (AGAP006765), *An. gambiae* sNPF (A0SIF1), *Cu. quinquefasciatus* NPF (JX317645.1), *Cu. quinquefasciatus* RYa (CPIJ008988), *Cu. quinquefasciatus* sNPF (CPIJ009049), *D. melanogaster* NPF (CG10342), *D. melanogaster RYa* (CG40733), *D. melanogaster* sNPF (CG13968). Receptor genes (accession numbers): *Ae. aegypti* NPYLR1 (U5N0U1), *Ae. aegypti* NPYLR5 (U5N1G6), *Ae. aegypti* NPYLR7 (U5N1H5), *Ae. aegypti* NPYLR8 (U5N0V1), *Ae. aegypti* PDFR (AAEL009024), *Ae. albopictus* sNPFR (AALF002670), *Ae. albopictus* RYaR1 (AALF021539), *Ae. albopictus* RYaR2 (AALF003651), *Ae. albopictus* NPFR1 (AALF023252), *Ae. albopictus* NPFR2 (AALF007614), *An. gambiae* sNPFR (A0SIF2), *An. gambiae* RYaR1 (AGAP000351), *An. gambiae* RYaR2 (AGAP000115), *An. gambiae* NPFR1 (AGAP004122), *An. gambiae* NPFR2 (AGAP004123), *C. quinquefasciatus* sNPFR (CPIJ013069), *C. quinquefasciatus* RYaR1 (CPIJ019394), *C. quinquefasciatus* RYaR2 (CPIJ018504), *C. quinquefasciatus* NPFR1 (CPIJ018265), *C. quinquefasciatus* NPFR2 (CPIJ006984), *D. melanogaster* sNPFR (CG7395), *D. melanogaster* RYaR (CG5811), *D. melanogaster* NPFR (CG1147).

### Peptide synthesis

Mature *Ae. aegypti* HP-I (pERPhPSLKTRFa), *Ae. aegypti* HP-III (pERPPSLKTRFa), and *Ae. aegypti* HP-I [Cys10] (pERPh PSLKTRC) were synthesized by The Rockefeller University Proteomics Resource Center. *Ae. albopictus* HP-I peptides (pERPhPSLKTRWa and pERPhPSYKLRFa), *Ae. aegypti* sNPF-3 (APSQRLRWa) and *Ae. aegypti* PDF (NSELNSLLSLPKKLNDAa) were synthesized by Bachem. Two stable isotope versions of *Ae. aegypti* HP-I were synthesized by The Rockefeller University Proteomics Resource Center: pERPhPS(^13^C_6_ ^15^N_1_ LKTRFa (+7 Da HP-I “medium”) and pERP(Arg-13C_6_ ^15^N_1_)hPS(^13^C_6_ ^15^N_1_ LKTRFa (+13 Da HP-I “heavy”).

### Neuropeptide receptor cloning

*Ae. aegypti* NPYLR-expressing plasmids were previously described (21). Full-length cDNAs for *An. gambiae*, *Ae. albopictus*, and *Cu. quinquefasciatus* receptors were synthesized by GenScript and subcloned with XhoI-NotI into the pME18s vector for expression in mammalian cells. Gene names (accession numbers) for the receptors used in this study are: *An. gambiae* RYaR2 (AGAP000115), *Ae. aegypti* NPYLR7 (AAEL008296), *Cu. quinquefasciatus* RYaR2 (CPIJ018504), *D. melanogaster* RYaR (CG5811), *An. gambiae* RYaR1 (AGAP000351), *Cu. quinquefasciatus* RYaR1 (CPIJ01934), *Ae. aegypti* NPYLR5 (AAEL017049), *Ae. albopictus* RYaR1 (AALF021539), *Ae. albopictus* RYa2 (AALF003651), *D. melanogaster* sNPFR (CG7395), *An. gambiae* sNPFR (AGAP012378, A0SIF2), *Cu. quinquefasciatus* sNPFR (CPIJ013069), *Ae. albopictus* sNPFR (AALF002670), *Ae. aegypti* NPYLR1 (AAEL013505), *D. melanogaster* NPFR (CG1147), *An. gambiae* NPFR1 (AGAP004122), *An. gambiae* NPFR2 (AGAP004123), *Cu. quinquefasciatus* NPFR1 (CPIJ018265), *Cu. quinquefasciatus* NPFR2 (CPIJ006984), *Ae. aegypti* NPYLR8 (AAEL010626), *Ae. albopictus* NPFR1 (AALF023252), *Ae. albopictus* NPFR2 (AALF007614). In three cases, we encountered receptor genes with separate annotation entries that encode highly similar (*Ae. albopictus* RYaR1 S31P and P32 L) or identical (*A.gambiae* NPFR1 and 2 and *Cu. quinquefasciatus* NPFR1 and NPFR2) proteins. In these cases, we selected one for expression analysis. Genes (accession numbers): *Ae. albopictus* RYaR1 (AALF021539) [not RYa2 (AALF003651)], *An. gambiae* NPFR2 (AGAP004123) [not NPYR1 (AGAP004122)], *Cu. quinquefasciatus* NPFR1 (CPIJ006984) [not NPFR2 (CPIJ018265)].

### Cell-based assays

HEK-293T cells were maintained using standard protocols in a Thermo Scientific FORMA Series II – Water Jacketed CO_2_ incubator. Cells were transiently transfected with 1 μg each of plasmid expressing GCaMP6s, Gqα15, and a test receptor using Lipofectamine 2000 (Invitrogen). Transfected cells were seeded into 384 well plates, and incubated overnight in DMEM media supplemented with Fetal Bovine Serum (Invitrogen) at 37**°**C and 5% CO_2_. Cells were imaged in reading buffer [Hanks’s Balanced Salt Solution (GIBCO) + 20 mM HEPES (Sigma-Aldrich)] using GFP-channel fluorescence of a Hamamatsu FDSS-6000 kinetic plate reader at The Rockefeller University High-Throughput Screening Resource Center. Compounds were prepared at 3x concentration in reading buffer in a 384-well plate (Greiner Bio-one). Plates were imaged every 1 sec for 5 min. 10 μl of compound was added to each well containing cells in 20 μl of reading buffer after 30 sec of baseline fluorescence recording. Fluorescence was normalized to baseline, and responses were calculated as max ratio (maximum fluorescence level/baseline fluorescence level).

### HP-I mutant generation

The HP-I gene was mutated using CRISPR-Cas9 methods as previously described (22). In brief, a 23 nucleotide guide RNA was designed to target the *HP-I* gene (target sequence with PAM underlined: AAAGACACGTTTCGGACGTT<u>CGG</u>). Purified guide RNA (25 ng/μl), Cas9 mRNA (300 ng/μl), and a DNA plasmid containing a homologous recombination sequence including a fluorescent marker (700 ng/μl) were injected into 1,131 pre-blastoderm stage *Ae. aegypti* embryos (Orlando strain) by the University of Maryland Insect Transformation Facility. 222 G0 animals survived, for a final hatch rate of 19.6%. G0 pupae were sexed and separated into male and female groups prior to eclosion. Male and female G0 adults were outcrossed to wild-type Orlando animals in batches of 20 G0 and 20 wild-type mates. F1 animals were screened for fluorescence to detect insertion of the fluorescent marker, but none were recovered. We therefore screened F1 animals for insertions or deletions at the *HP-I* locus. 174 F1 adults were intercrossed in groups of 3 females and 3 males and analyzed with Illumina MiSeq for insertions/deletions surrounding the cut site. Animals were pooled into groups of 3 for genomic DNA extractions and MiSeq amplicon generation. PCR primers used to generate MiSeq amplicons (LD15F CGAGGATCAACGTTAGTGTCATATA; LD15R GAGCCGAGCGCTTTTCCATTATGTC) generating a 170bp wild-type product. *HP-IΔ54* mutation was selected for isolation due to isolation of stable lines from both male and female founders and ease of genotyping. The *HP-IΔ54* mutant was genotyped by generating PCR products using the following primers: Forward (5’-CGTTAGTGTCATATAGTTGATTTTT-3’), Reverse (5’-TACTGACTCTGAGCCGAGCGCTTTT-3’). These products were readily discriminable on a 2% agarose gel (wild-type: 170 bp vs mutant: 116 bp). Genotypes were confirmed by Sanger DNA sequencing (Genewiz). Mutants were blood-fed on human subjects until a stable line was generated, and subsequently maintained by blood-feeding on mice.

### Liquid chromatography and mass spectrometry

Targeted LC-MS/MS was used to analyze HP-I in 7-10 day-old wild-type or *HP-IΔ54* mutant mosquitoes. Virgin males and females were obtained by sexing animals as pupae and housing them exclusively with same-sex siblings until proteomic sample preparation. Non-virgin males were group-housed with female siblings from eclosion until proteomic sample preparation. In mating experiments, 10 virgin females who had never taken a blood-meal were exposed in bucket cages at 25-28°C, 70-80% relative humidity to 11 sexually mature males for 10 min, 2 hr, or 24 hr. Males were removed, and females were immediately processed for proteomic analysis.

Whole animals 7-10 days post-eclosion were boiled in groups of 7-10 for 5 min at 100°C in 150 μl of MilliQ water. The water fraction was decanted into a separate tube and set aside. Extraction solution (150 μl 0.25% acetic acid) was added to the carcasses along with two stable isotope versions of HP-I (1 ng/mosquito of HP-I “medium” and 10 ng/mosquito of HP-I “heavy”), and tissue was homogenized using a Kontes pellet pestle grinder (Sigma-Aldrich). The water and acid fractions were centrifuged separately at 4°C for 30 min, and then supernatants combined and passed through a Microcon 10-kDa-molecular weight cutoff filter (Millipore, Merck KGaA) by centrifuging at 4°C for 30 min. Samples were spun to dryness in an Eppendorf Speedvac and resuspended in 20 μL 0.1% TFA/2% acetonitrile. 9 μL of each sample were separated by reversed phase (Acclaim 120 C18, 3um, 120A 2.1mm x 150mm, Thermo Fisher) coupled to an Orbitrap XL (Thermo Fisher Scientific) operated in positive mode. MS spectra were acquired at a resolution of 60,000@m/z 400 and the triply charged endogenous mature HP-I (m/z 409.9034), in addition to +7 Da (HP-I “medium”) (m/z 412.2425) and +13 Da (HP-I “heavy”) (m/z 414.2471) stable isotope versions of HP-I, was continually targeted by MS/MS and measured in the ion trap. Peptides were isolated using a window of 2.0 m/z. Peptides were eluted at 200 μL/min, increasing from 7% Buffer B/93% Buffer A to 35% Buffer B/65% Buffer A over a period of 13 min (Buffer A: 0.1% formic acid; Buffer B: 0.1% formic acid in acetonitrile). Between each sample, the column was cleaned for 3 minutes in 90% Buffer B /10% Buffer A. The column was then conditioned for 5 minutes with 100% Buffer A. All solvents were HPLC grade. Both MS and MS/MS signals were extracted and analyzed using Skyline (40). To estimate recovery, signals of the spiked-in stable isotope-labeled HP-I peptide were compared to a dilution series of measurements of known amount of the stable isotope-labeled HP-I peptide.

### Glytube blood-meal feeding

For experiments in Fig. 2 to Fig. 4, females were fed sheep blood using Glytube membrane feeders exactly as described (36). Glytubes were placed on top of mesh on the mosquito cage, and females were allowed to feed through the mesh for 15 min. Fed females were scored by eye for engorgement of the abdomen and weighed to confirm feeding status. Females scored as partially fed were discarded.

### Egg-laying assays

7 to 14 day-old female mosquitoes were fed sheep blood using Glytube membrane feeders (36). Immediately after blood-feeding, individual mosquitoes were placed in plastic *Drosophila* vials (25 mm diameter, 95 mm long) containing 5 ml water and a Whatman filter paper (55 mm diameter; GE Healthcare) folded into a cone to act as an oviposition substrate. At 144 hr post-blood-meal, filter papers were removed, and eggs were manually counted by eye.

### Uniport olfactometer

Host-seeking behavior was measured using a uniport olfactometer exactly as described (21).

### *ECFP* paternity marker strain generation

A genetically modified strain generated for an unrelated study (C.J. McMeniman, in preparation) was utilized as a paternity marker because it expressed high levels of ubiquitous ECFP in larvae, had normal capacity to produce offspring (Fig. 3A), and showed wild-type levels of paternity enforcement (Fig. 3D and E). A zinc-finger nuclease (ZFN) targeting *Ae. aegypti* AAEL002167 was produced by the CompoZr Custom ZFN Service (Sigma-Aldrich Life Science). The nucleotide sequence of the ZFN binding and wild-type heterodimeric Fok1 endonuclease sites for this ZFN pair are denoted in upper case and lower case letters, respectively: 5’-CCACACTTCTGGATTCCATtcgtaGGATGGGGAGTAGCA-3’. Homologous recombination was used to insert a poly-Ubiquitin-ECFP cassette (pSL1180-HR-*PUb*ECFP, Addgene plasmid #47917) into the locus. Full details of strain generation are available upon request from C.J.M.

### NPYLR1 mutant strain

Experiments using *NPYLR1* mutants used in this study carried an 8 bp deletion (*NPYLR1Δ8*) as previously described (21). Strains were genotyped prior to use to confirm the presence of the allele in a homozygous state (21).

### Mating and paternity assays

Mosquitoes were separated by sex at the pupal stage and sex was confirmed within 24 hr of eclosion. Females were separated into small groups (n = 7-10) and housed in 473 ml paper soup cups (Webstaurant Store) overnight 25-28°C, 70-80% relative humidity. 6 females were exposed to 7 males for the indicated times, using an aspirator (John W. Hock Company) to introduce and remove males from the soup cups. For remating experiments, male 2 was exposed to the female for 30 min at 25-28°C, 70-80% relative humidity. Immediately after assay termination, animals were anesthetized at 4°C and separated by sex. Females were allowed to recover overnight at 25-28°C, 70-80% relative humidity, blood-fed, and offered an oviposition substrate in a vial or a cup to lay eggs. In remating assays, larvae were screened for ECFP fluorescence 3-4 days after hatching by transferring them to a wet filter paper, counting the total number of larvae manually by eye, and scoring the number of ECFP-positive animals using a CFP filter on a Nikon SMZ-1500 upright microscope.

### Peptide Injections

Twelve ∼12-day-old female mosquitoes of each genotype were anesthetized at 4^o^C for 30 min, and placed on an acrylic grid for injection at 4°C. Animals were injected in the thorax with 150 nl of each solution using a Drummond Nanoject II (Catalogue #3-000-204) attached to 3.5″ pipettes (Drummond, catalogue #3-000-203-G/X) pulled on a micropipette puller (Sutter Instruments Co., Model P-97). Peptides were dissolved at 500 μM in buffer (1X Ca^+2^/Mg^+2^-free PBS; Lonza, catalogue #17517Q). Mosquitoes were allowed to recover in groups for 12-16 hours at 25-28°C, 70-80% relative humidity with access to water, and then mated and blood-fed as described above. Eggs were collected, hatched individually and scored by eye for the presence of larvae 3-4 days after hatching. There was no difference in the number of viable offspring produced by uninjected wild-type and *NPYLR1* females (Mann-Whitney test; see Data file 1).

### Data Analysis

All statistical analysis was performed using Prism (Graphpad Software).

